# Determinants and Mechanisms of the Low Fusogenicity and Endosomal Entry of Omicron Subvariants

**DOI:** 10.1101/2022.10.15.512322

**Authors:** Panke Qu, John P. Evans, Chaitanya Kurhade, Cong Zeng, Yi-Min Zheng, Kai Xu, Pei-Yong Shi, Xuping Xie, Shan-Lu Liu

## Abstract

The rapid spread and strong immune evasion of the SARS-CoV-2 Omicron subvariants has raised serious concerns for the global COVID-19 pandemic. These new variants exhibit reduced fusogenicity and increased endosomal entry pathway utilization compared to the ancestral D614G variant, the underlying mechanisms of which remain elusive. Here we show that the C-terminal S1 mutations of the BA.1.1 subvariant, H655Y and T547K, critically govern the low fusogenicity of Omicron. Notably, H655Y also dictates the enhanced endosome entry pathway utilization. Mechanistically, T547K and H655Y likely stabilize the spike trimer conformation, as shown by increased molecular interactions in structural modeling as well as reduced S1 shedding. Importantly, the H655Y mutation also determines the low fusogenicity and high dependence on the endosomal entry pathway of other Omicron subvariants, including BA.2, BA.2.12.1, BA.4/5 and BA.2.75. These results uncover mechanisms governing Omicron subvariant entry and provide insights into altered Omicron tissue tropism and pathogenesis.

## Introduction

The emergence and rapid spread of the Omicron subvariants of SARS-CoV-2 around the world has caused serious concern about vaccine efficacy because of the large numbers of mutations present in this lineage, with more than 30 substitutions in the Spike (S) proteins (Viana et al., 2022). Indeed, many recent studies have shown that Omicron is resistant to neutralization by antibodies induced by two-dose mRNA vaccination that is greatly restored by booster vaccination (Carreno et al., 2022; Evans et al., 2022; Liu et al., 2022a; Perez-Then et al., 2022; Planas et al., 2022; Xia et al., 2022; Yu et al., 2022). Consequently, Omicron led to a global surge of COVID-19 cases. BA.1.1 and BA.1 were responsible for the initial Omicron wave, but were replaced by the BA.2 subvariant, which was more transmissible and caused reinfection in patients who previously were infected by BA.1 (Marc Stegger et al., 2022). Remarkably, derivatives of BA.2, including BA.2.12.1 which subsequently became predominant in the US, and the BA.4 and BA.5 subvariants bearing identical S proteins (referred to as BA.4/5 hereafter), which are currently dominant in the world, are driving a new surge in cases as both have stronger immune escape ability, especially the BA.4/5 variants (Cao et al., 2022; Hachmann et al., 2022; Qu et al., 2022a; Wang et al., 2022). Despite the gradually improved virus transmissibility observed in these subvariants, they appear to cause milder disease compared to Delta and other previous variants (Diamond et al., 2021; Halfmann et al., 2022; McMahan et al., 2022). The mechanism underlying the reduced pathogenicity of Omicron is currently unclear but has been under intense investigation.

The SARS-CoV-2 S protein is a typical class I viral fusion protein that utilizes the cognate receptor angiotensin-converting enzyme 2 (ACE2) for binding and cellular entry (Hoffmann et al., 2020; Zhou et al., 2020). It has a furin cleavage site at the S1/S2 junction of S, which facilitates SARS-CoV-2 entry at the plasma membrane, viral replication in human lung epithelial cells, as well as transmission in animals (Mykytyn et al., 2021; Peacock et al., 2021). As SARS-CoV-2 evolves, its S has gained a large number of mutations and possibly undergone recombination, resulting in the emergence of the initial D614G variant and several major variants of concern (VOCs) such as Alpha, Beta, Gamma, Delta, and Omicron, some of which directly or indirectly increase S cleavage to enhance fusogenicity (Escalera et al., 2022; Liu et al., 2022b; Saito et al., 2022). Intriguingly, results from many groups including ours have shown that the Omicron subvariants exhibit substantially impaired cell-cell fusion capacity and tend to use the endosomal entry pathway mediated by cathepsin L/B (Cat L/B), rather than the plasma membrane entry pathway mediated by transmembrane protease serine 2 (TMPRSS2) which is preferred by other previous variants (Du et al., 2022; Meng et al., 2022; Qu et al., 2022b).

Currently, the underlying molecular mechanism by which Omicron subvariants use endosomal entry more efficiently than plasma membrane entry is unclear. Herein, we provide evidence that some key mutations in S1 and/or the S1/S2 junction region of BA.1.1 S, such as T547K and H655Y, especially the latter, dictate its intrinsically low fusogenicity and endosomal entry. Mechanically, we find that T547K and H655Y restrict BA.1.1 S-mediated cell-cell fusion possibly through stabilizing the S trimer conformation. Remarkably, H655Y, but not T547K which is only carried by BA.1.1 and BA.1 of Omicron subvariants (Qu *et al.*, 2022a), governs the entry preference and fusion capability of all predominant Omicron subvariants. Together, our results reveal that mutations in S1 around the furin cleavage site of Omicron S critically modulate the unique biology of Omicron entry and are potentially associated with pathogenesis.

## Results

### Critical amino acid residues dictating the differential entry patterns of Omicron subvariant BA.1.1 in HEK293T-ACE2 and Calu-3 cells

Previous studies have shown that SARS-CoV-2 entry is cell-type dependent, depending on the efficiency of furin cleavage of the virus S protein and also the levels of TMPRSS2 expression on the plasma membrane of target cells (Bestle et al., 2020; Peacock *et al.*, 2021). In low TMPRSS2-expressing 293T-ACE2 and Vero cells, the endosomal entry pathway is predominant; however, in Calu-3 and other cells expressing high levels of TMPRRS2, SARS-CoV-2 primarily uses the plasma membrane for entry (Hoffmann *et al.*, 2020). We hypothesized that mutations at the C-terminus of S1, or near the furin cleavage site of Omicron S, potentially govern its endosomal entry. To test this hypothesis, we created reversion mutations specific to residues T547K, H655Y, N679K, and P681H of the Omicron subvariant BA.1.1 (**Fig. 1a**), and examined their impact on the entry of BA.1.1 into HEK293T-ACE2, HEK293T-ACE2-TMPRSS2 and Calu-3 cells using our previously reported HIV lentiviral pseudotyping system (Zeng et al., 2020). In parallel, the entry in these cell types was tested for the corresponding forward mutations, i.e., T547K, H655Y, N679K, and P681H made in the backbone of the ancestral D614G construct (**Fig. 1a**). We found that, compared with parental BA.1.1, the reversion mutation Y655H exhibited a profound reduction of entry efficiency in HEK293T-ACE2 (**Fig. 1b**), and in HEK93T-ACE2-TMPRSS2 cells, albeit to a lesser extent (**Fig. 1c**), yet apparently enhanced entry in Calu-3 cells (**Fig. 1d**), suggesting that H655Y is the most critical change in BA.1.1 S that distinguishes its entry in these different cell types. Consistent with this, we observed that the forward mutation H655Y exhibited reduced entry efficiency in Calu-3 cells, but increased entry in 293T-ACE2 and 293T-ACE-TMPRRS2 cells (**Figs. 1e-f**). While similar results were also observed for the BA.1.1 K547T mutation, the impact on entry was not as dramatic as that of H655Y (**Figs. 1b-g**). Surprisingly, two other mutations of BA.1.1, N679K and P681H, which are in close proximity to the furin cleavage site, did not seem to obviously affect viral entry in these cell types (**Figs. 1b-g**).

**Figure 1:**
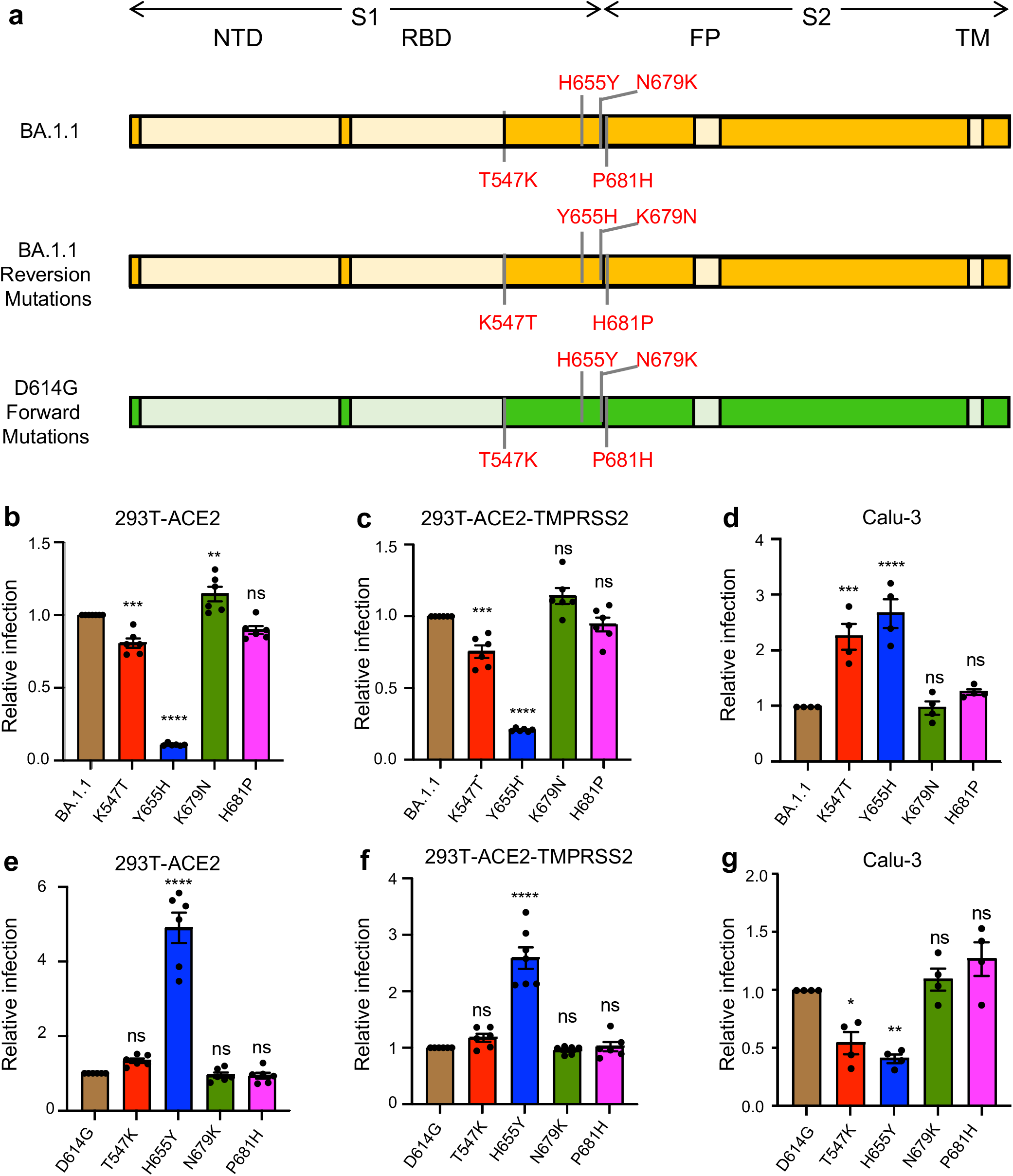
H655Y governs differential entry of Omicron BA.1.1 into distinct target cells. **(a)** Schematic representations of BA.1.1 and D614G Spike glycoprotein are presented. The N-terminal domain (NTD), the receptor binding domain (RBD), the fusion peptide (FP) and the transmembrane (TM) region are indicated. Only the mutations at the C-terminus of S1 and those near the S1/S2 junction in BA.1.1 S relative to SARS-CoV-2 D614G S are shown (**Top**, BA.1.1) (Mlcochova et al.); also displayed are these four amino acid mutations of BA.1.1 S replaced with the corresponding amino acid in the D614G S (**Middle**, BA.1.1 reversion mutations), and these four residues in D614G S substituted with the corresponding amino acid in BA.1.1 S (**Bottom**, D614G forward mutations). (**b-d)** The relative infectivity of pseudotyped viruses encoding BA.1.1 S with single mutated amino acids replaced with the corresponding amino acid in the D614G S in HEK293T-ACE2 cells (n=6) (**b**), HEK293T-ACE2-TMPRSS2 cells (n=6) (**c**), and Calu-3 cells (n=4) (**d**). The luciferase activity of parental BA.1.1 was set as 1.0 for comparison. (Deng et al.) The relative infectivity of pseudotyped viruses encoding D614G S with single amino acids substituted with the corresponding amino acid in the BA.1.1 S in HEK293T-ACE2 cells (n=6) (**e**), HEK293T-ACE2-TMPRSS2 cells (n=6) (**f**), and Calu-3 cells (n=4) (**g**). The luciferase activity of parental D614G was set as 1.0. In call cases, bars represent means +/− standard error, and significance was determined by one-way ANOVA with Bonferroni’s multiple testing correction. P-values are represented as ns indicates p ≥ 0.05, *p < 0.05, ***p < 0.001, ****p < 0.0001.

### H655Y governs the endosomal entry pathway of BA.1.1 – differential effects of E64d and Camostat

Previous studies have shown that SARS-CoV-2 entry in HEK293T-ACE2 cells is predominantly endosomal whereas in Calu-3 cells entry is primarily through the plasma membrane (Peacock *et al.*, 2021). We thus decided to use HEK293T-ACE2-TMPRSS2 cells, which allow entry through both endosomal and plasma membrane routs, to determine the impact of these BA.1.1 mutants on entry in the presence of the endosomal Cat L/B inhibitor E64d or the TMPRSS2 inhibitor Camostat. While BA.1.1 was more sensitive to treatment by E64d but less sensitive to treatment by Camostat compared to D614G, BA.1.1-Y655H was less sensitive to E64d than BA.1.1, with a half maximal inhibitory concentration (IC_50_) of > 25 μM versus an IC_50_ of 0.4 μM for the parental BA.1.1 (**Figs. 2a** and **2g**). Additionally, BA.1.1-Y655H was more sensitive to Camostat than BA.1.1, with an IC_50_ of 10.3 μM versus IC_50_ >150 μM for BA.1.1 (**Figs. 2b** and **2g**). Convincingly, the forward mutant D614G-H655Y was more sensitive to E64d, with an IC_50_ of 0.8 μM versus 23.5 μM for the parental D614G (**Figs. 2c** and **2g**), but less sensitive to Camostat, with an IC_50_ of 91.3 μM versus 11.6 μM for the parental D614G (**Figs. 2d** and **2g**). Overall, we discovered that the BA.1.1 reversion mutant Y655H had a more than 14.6-fold decreased IC_50_ for Camostat and an over 62.5-fold increased IC_50_ for E64d compared to the parental BA.1.1 (**Fig. 2g**). Further, the forward mutant H655Y has a 7.9-fold increased IC_50_ for Camostat and a 29.4-fold decreased IC_50_ for E64d relative to the parental D614G (**Fig. 2g**). Of note, the T547K mutation of BA.1.1 did not seem to have a dramatic impact on the sensitivity to these drug treatments (**Figs. 2a-d,** and **2g**).

**Figure 2:**
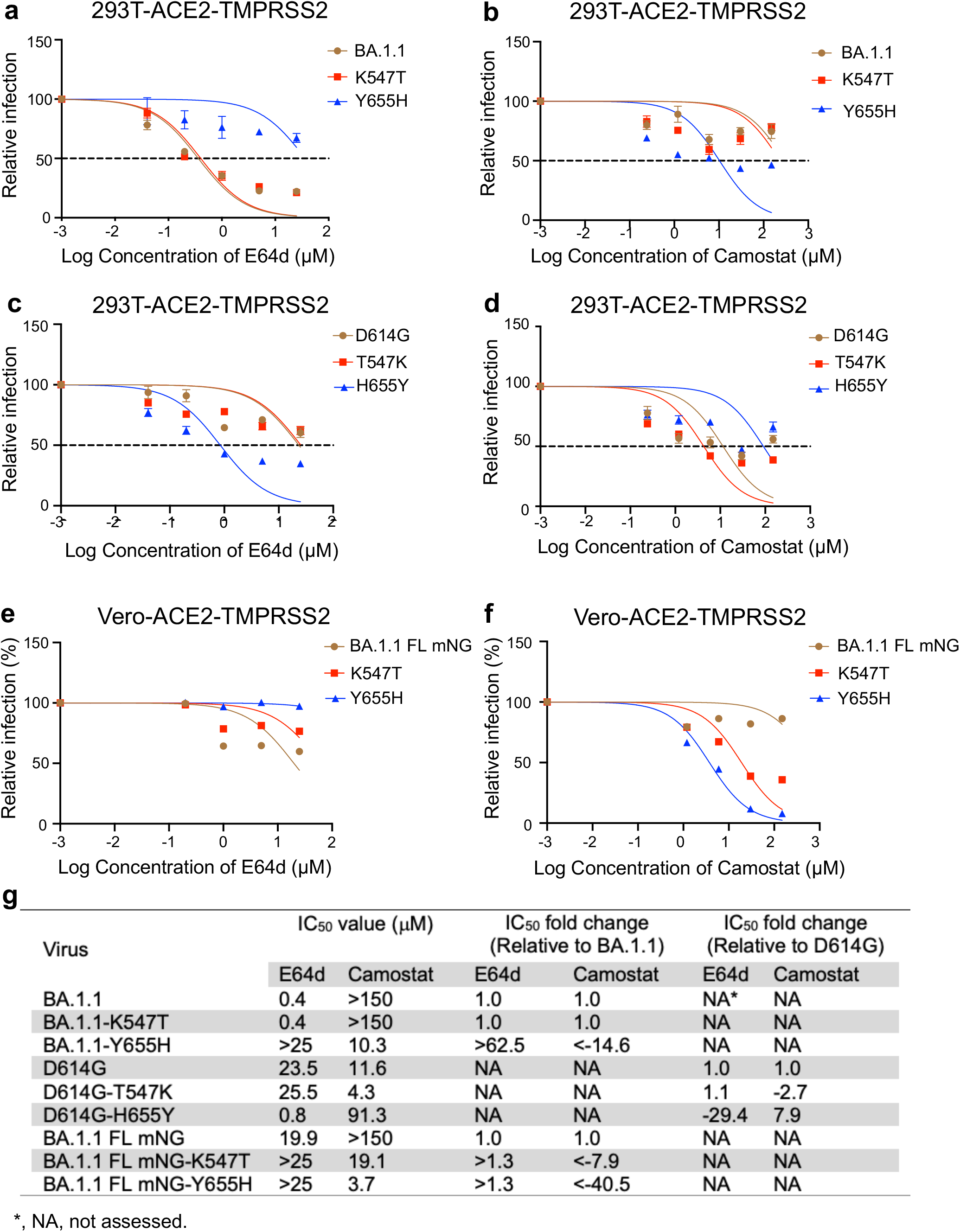
H655Y dictates BA.1.1 entry through the endosomal pathway. HEK293T-ACE2-TMPRSS2 cells were pretreated with E64d (**a** and **c**) or Camostat (**b** and **d**), and then transduced with the indicated pseudotyped viruses in the presence of varying concentrations of the inhibitors. The relative infection was calculated by setting the luciferase activity of each virus at the 0 μM of the inhibitors as 100%, and the half maximal inhibitory concentration (IC_50_) values were determined by non-linear regression with least squares fit. Vero-ACE2-TMPRSS2 cells were pretreated with E64d (**e**) or Camostat (**f**), followed by infection with parental BA.1 FL mNG or its mutants. The infection ratio was determined by flow cytometry, and relative infection was calculated by setting the infection percentage of each virus in the absence of the drugs as 100%. (**g**) IC_50_ values of each virus in the presence of the individual drugs are shown. Additionally, the IC_50_ fold change relative to parental BA.1 (or BA.1 FL mNG) or D614G are displayed.

To confirm the above pseudotype lentivirus results in an authentic SARS-CoV-2 system, we engineered the complete BA.1 full length (FL) spike (Pitts et al., 2022) into the infectious cDNA clone of USA-WA1/2020 with mNeonGreen (Viana *et al.*) reporter (Xie et al., 2020) to obtain BA.1-FL mNG SARS-CoV-2 and then generated the individual reversion mutations of spike K547T or Y655H in this background. Using these infectious viruses, we determined the impact of these two mutations on BA.1 entry into Vero-ACE2-TMPRSS2 cells using the inhibitors E64d or Camostat. The results showed that the parental BA.1-FL mNG SARS-CoV-2 was the most sensitive to E64d with an IC_50_ of 19.9 μM among the three viruses, followed by BA.1-FLmNG-K547T and then BA.1-FL mNG-Y655H, although these variants were generally much less sensitive to E64d compared to that in 293T-ACE2 cells, with no exact IC_50_ value calculated for the two mutants (**Figs. 2e**, **2g and S1a**). In contrast, compared to the parental BA.1-FL mNG which was resistant to Camostat treatment with an IC_50_ of more than 150 μM, BA.1-FL mNG-K547T was sensitive to Camostat with an IC_50_ of 19.1 μM, a more than 7.9-fold reduction in IC_50_ relative to BA.1-FL mNG; BA.1-FL mNG-Y655H was more sensitive to Camostat, with an IC_50_ of 3.7 μM, approximately > 40.5-fold decreased IC_50_ relative to BA.1.1 FL mNG (**Figs. 2f, 2g and S1b**). These results together indicated T547K and H655Y mutations, especially the latter, greatly contribute to the altered entry preference of BA.1 in target cells. Overall, the results of pseudotyped as well as authentic viruses highlight the essential of role of H655Y mutation in dictating the Omicron BA.1 subvariant S-mediated endosomal entry compared to the ancestral D614G, which enters target cells predominantly through the plasma membrane.

### H655Y critically controls the low fusogenicity of BA.1.1 Spike

To investigate the underlying mechanism of the T547K and H655Y mutations in modulating BA.1.1 entry preference, we examined the role of these two mutants along with N679K, P681H and parental BA.1.1 in S expression and S-mediated membrane fusion. In parallel, the impacts of the corresponding forward mutants were also investigated. Flow cytometric analysis of the S surface expression in HEK293T cells used to produce pseudotyped lentivirus showed that all these reversion mutants had approximately comparable levels of expression to each other and the parental BA.1.1, except for the reversion mutant K547T, which had modestly reduced surface expression (**Figs. 3a-b**), while all the forward mutants had similar levels of expression to D614G (**Fig. S2a-b**). We then conducted syncytia formation assays in HEK293T-ACE2 cells transfected to express GFP and the individual S constructs and observed that K547T and Y655H substantially promoted the S-mediated cell-cell fusion, whereas K679N and H681P had no effect (**Figs. 3c-d**). Surprisingly, while forward mutations H655Y and T547K slightly reduced the D614G S-induced syncytia, two other mutations N679K and P681H enhanced D614G S-mediated fusion (**Figs. S2c-d**), similar to some previous reports (Rajah et al., 2021).

**Figure 3:**
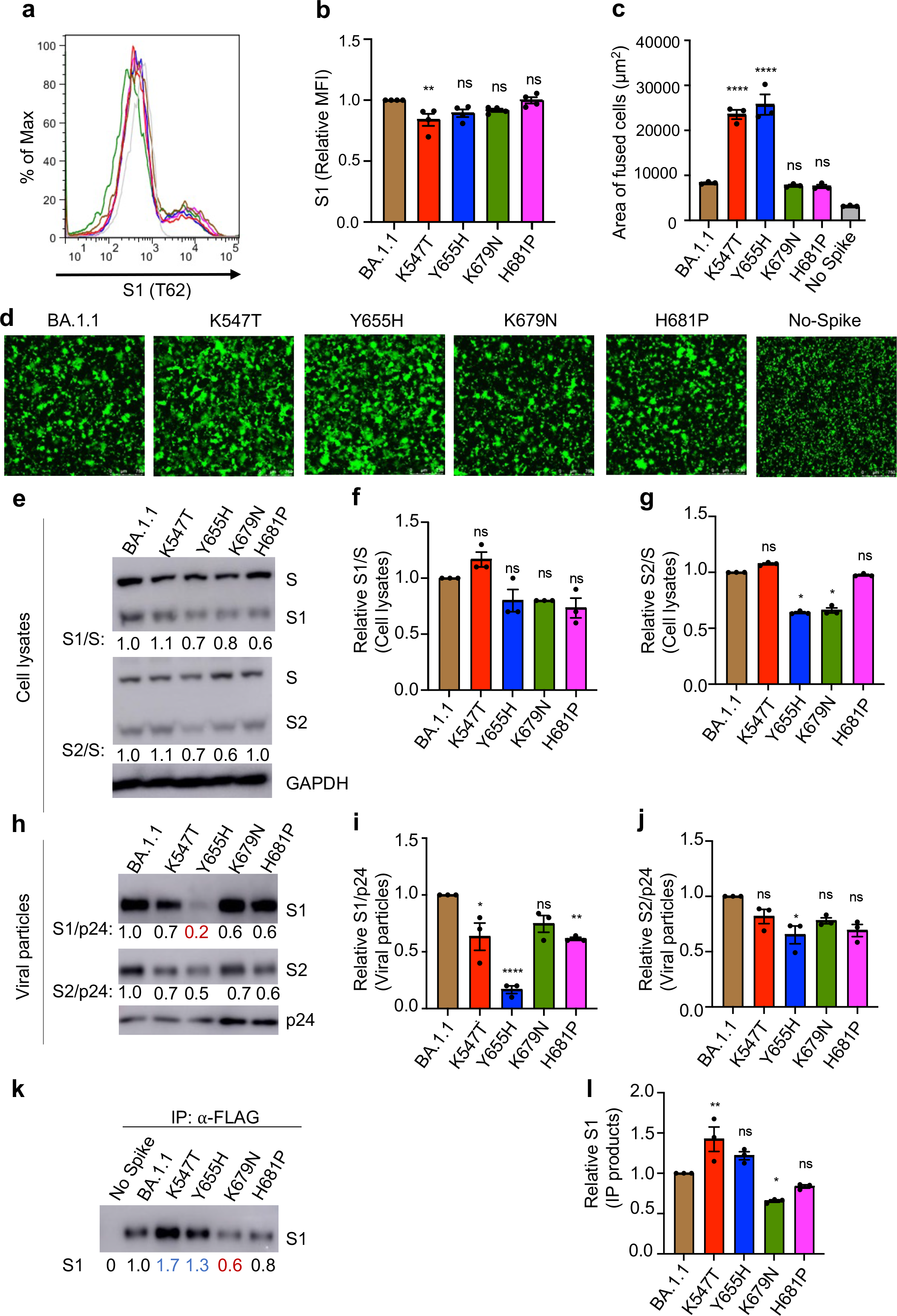
H655Y critically controls the low fusogenicity of BA.1.1 S. (**a**) The S expression on the cell surface of HEK293T cell used to produce pseudotyped lentivirus was determined by staining cells with anti-SARS-CoV-2 S1 (T62) antibodies. Representative flow cytometric histograms of the S1 signal are shown. (**b**) Mean fluorescence intensity of S1 signal was calculated and shown, n=4. (**c-d**) Syncytia formation was assayed in HEK293T-ACE2 cells transfected to express S construct of interest and GFP. The average area of fused cells from one of five independent experiments (**c**) and representative syncytia images (**d**) are shown. Scale bars represent 150 μm. (**e**) Producer cell lysates were probed for S1 subunit, S2 subunit, p24, and GAPDH, respectively. S signals were quantified using NIH ImageJ and relative efficiency of S cleavage was calculated by setting the ratio of S1/S or S2/S of BA.1.1 to 1.00. Representative blots are shown. The relative ratios of S1/S (**f**) and S2/S (**g**) from three independent experiments are shown. (**h)** The pseudovirions were purified and blotted for S1, S2, and p24, and S1 or S2 signals in virions were quantified and normalized to p24. The relative S1/p24 or S2/p24 was calculated by setting the ratio of BA.1.1 to 1.00. The relative S1/p24 (**i**) and the relative S2/p24 (**j**) ratios from three independent experiments are shown. (**k-l)** Reversion of H655Y to Y655H enhances BA.1.1 S1 shedding. HEK293T cells transfected with indicated S constructs were treated with soluble ACE2-Fc (sACE2), followed by immunoprecipitation (IP) of S1 in the culture media by anti-FLAG beads. S in the IP products (**k**) were probed for SARS-CoV-2 S1, with GAPDH as a loading control. Relative S1 in the IP products is shown (**l**), calculated by setting the S1 signals of BA.1.1 to 1.0. The error bars in (**b**, **c**, **f**, **g**, **i**, **j** and **l)** are means +/− standard error; ns indicates p ≥ 0.05, *p < 0.05, **p < 0.01, ****p < 0.0001.

### Reversion of H655Y to Y655H results in increased BA.1.1 S1 shedding

The efficiency of SARS-CoV-2 S cleavage by furin is known to be associated with membrane fusogenicity (Johnson et al., 2021; Saito *et al.*, 2022). We next examined how these mutations might influence BA.1.1 S processing by analyzing the ratio of S1/S and S2/S in viral producer cells and purified viral particles. In cell lysates, while K547T exhibited a slightly increased or comparable ratio of S1/S and S2/S, H681P exhibited a decreased S1/S ratio but not S2/S ratio, Y655H and K679N showed a decreased ratio of both S1/S and S2/S (**Figs. 3e-g**). However, none of the four forward mutants showed obvious change in S1/S1 or S2/S ratios in the cell lysates, except for N679K which exhibited a modestly increased S2/S ratio (**Figs. S3a-c**), suggesting that these four mutations do not appear to significantly modulate BA.1.1 furin cleavage in viral producer cells. Interestingly, we found that all four reversion mutants, most notably Y655H, showed a decreased level of S1 and S2 compared to the parental BA.1.1 in viral particles after being normalized by p24 (**Figs. 3h-j**). Additionally, the corresponding forward mutations T547K and H655Y, especially the latter, substantially increased the S1/p24 and S2/p24 ratios in viral particles relative to D614G (**Figs. S3d-f**). The drastically decreased level of S1 for Y655H in viral particles corresponded to its significantly reduced infectivity in HEK293T-ACE2 and HEK293T-ACE2-TMPRSS2 cells, potentially due to premature inactivation by high level expression of ACE2 in these cells, despite apparently increased infectivity in Calu-3 cells (**Figs. 1b-d**). These results, along with the increased fusogenicity of Y655H, suggest that the H655Y mutation in BA.1.1 critically governs its low fusion activity and differential entry between HEK293T-ACE2 and Calu-3 cells.

The comparable or modestly increased efficiency of furin cleavage in viral producer cells of K547T and Y655H reversion mutants, in contrast to their decreased levels of S1 in the virions, especially Y655H, were indeed surprising, especially given their significantly increased cell-cell fusion activity observed in HEK293T-ACE2 cells. One possibility is that the S1 subunit of these two reversion mutants, especially Y655H, was shed into culture media during virus production. Indeed, we found that, compared to the parental BA.1.1, K547T and Y655H exhibited increased levels of S1 shedding into culture media in the presence of treatment by soluble ACE2 (**Figs. 3k and l**). Of note, S1 shedding of two other reversion mutants, K679N and H681P, was decreased compared to the parental BA.1.1, which appeared to correlate with their relatively low furin cleavage (**Figs. 4k and l**). Altogether, these results suggest that the T547K and H655Y mutations in BA.1.1, especially the latter, stabilize the spike conformation, contributing to decreased fusogenicity and entry in Calu-3 cells.

**Figure 4:**
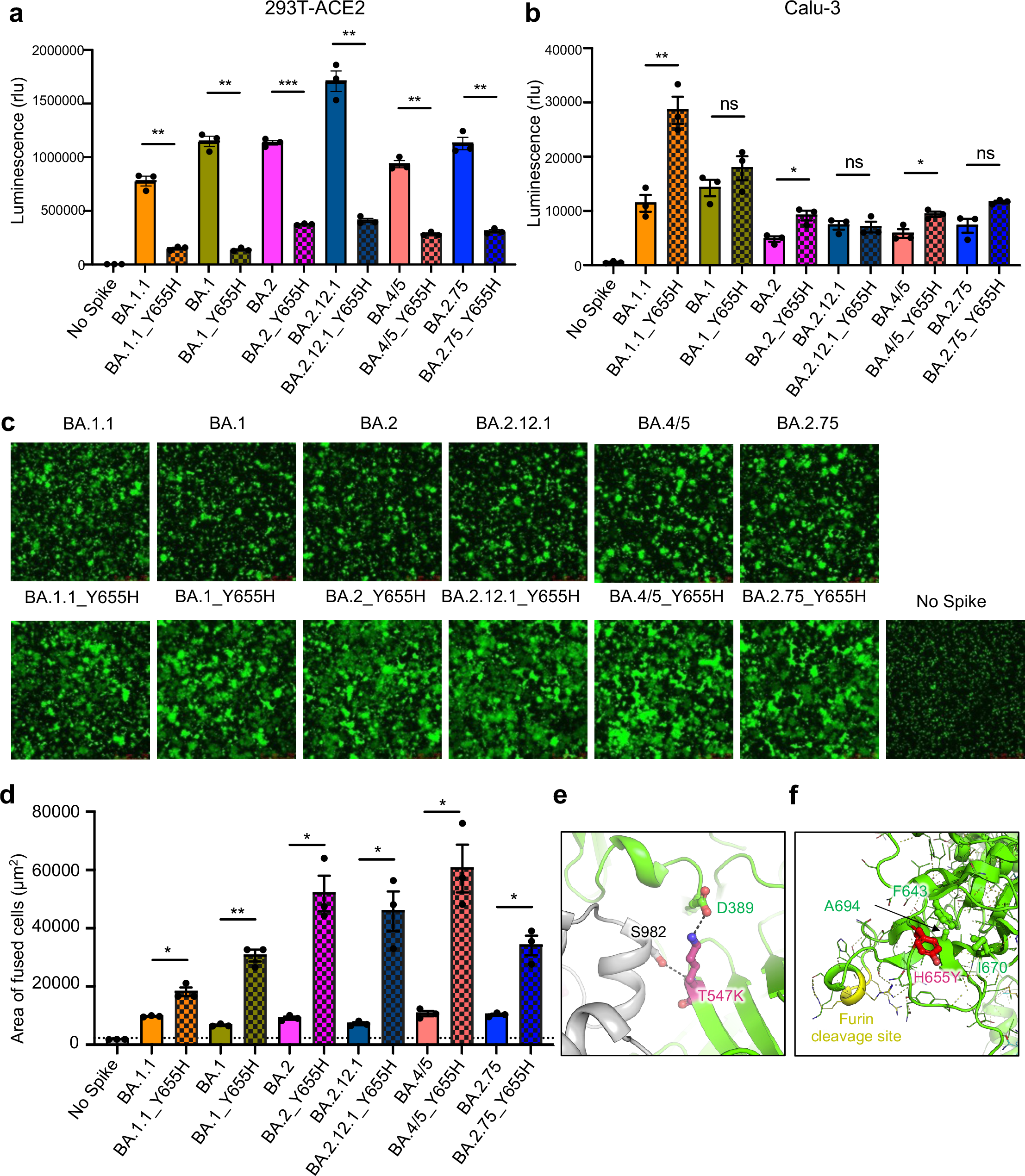
H655Y dictates the endosome entry preference and the low fusogenicity of all major Omicron subvariants. (**a**) Infectivity of pseudotyped viruses in HEK293T-ACE2 cells is shown, n=3. (**b**) Infectivity of pseudotyped lentivirus in CaLu-3 cells is also displayed, n=3. (**c-d**) Syncytia formation assays in HEK293T-ACE2 cells were performed as described in Fig. 3. Representative syncytia images (**c**) and the average area of fused cells (**d**) are shown, n=3. Scale bars represent 150 μm. (**e**) Molecular modelling. The T547K mutation introduces a salt-bridge interaction with the D389 located on the base of the RBD, and a hydrogen bond with residue S982 in the S2 domain. (**f**) BA.1.1 H655Y mutation reduces local conformational flexibility by creating hydrophobic contacts with residues F643, A694 and I670, which may interfere with the furin accessibility to the cleavage site nearby. Bars in (**a**), (**b**) and (**d**) represent means +/− standard error, and significance is determined by two-tailed Student’s t test with Welch’s correction. P-values are represented as ns for p ≥ 0.05, *p<0.05, **p < 0.01, ***p < 0.001.

### H655Y dictates the endosomal entry and low fusogenicity of other Omicron subvariants

As in BA.1.1, all previously or currently circulating Omicron subvariants contain the H655Y substitution, but not T547K (**Fig. S4**). We therefore predicted that the H655Y mutation also governs the entry preference and low fusogenicity of other variants. To this end, we introduced the H655Y reversion mutation into these Omicron subvariants, i.e., BA.1, BA.2, BA.2.12.1, BA.4/5 and BA.2.75, and examined their impact on entry of these subvariants in HEK293T-ACE2 and Calu-3 cells using the same HIV lentiviral pseudotyping system. We observed that the reversion mutation Y655H significantly reduced the entry efficiency of all these Omicron subvariants in HEK293T-ACE2 cells (**Fig. 4a**), but substantially promoted their entry in Calu-3 cells, although the increase in Calu-3 cells was generally modest or not seen in some cases (**Fig. 4b**). We also examined the influence of the reversion mutation Y655H on the fusogenicity of these Omicron subvariants using the syncytia formation assay described above. Again, we found that similar to BA.1.1, Y655H also significantly promoted the S-mediated syncytia formation of BA.1, BA.2, BA.2.12.1, BA.4/5, and BA.2.75 (**Figs. 4c-d**).

To provide molecular insight into how T547K and H655Y might dictate the low fusogenicity and endosomal entry of the Omicron subvariants, we performed molecular modeling of Omicron BA.1.1 spike protein. Indeed, we found that the T547K mutation could establish a salt-bridge interaction with the residue D389 located on the base of the receptor binding domain (RBD), as well as an inter-protomer hydrogen bond with the residue S982 on S2 domain, thus stabilizing its closed conformation and prevent S1 shedding (**Fig. 4e**). The mutation H655Y located in the proximity to the furin cleavage site in S could reduce local structural flexibility by forming hydrophobic interactions with residues F643, I670 and A694, which may interfere with or decrease the accessibility of furin to the cleavage site (**Fig. 4f**).

## Discussion

Omicron exhibits exceptional immune evasion (Garcia-Beltran et al., 2022; Kuhlmann et al., 2022; Planas *et al.*, 2022; Pulliam et al., 2022; Viana *et al.*, 2022) because of the presence of large numbers of mutations in the S protein. We and others have reported that Omicron also exhibits a distinct entry preference, substantially impaired furin cleavage, as well as decreased cell-cell fusion (Meng *et al.*, 2022; Qu *et al.*, 2022b; Zeng et al., 2021). However, the molecular mechanisms of these distinct features of the Omicron subvariants remain elusive. In this study, we interrogated key mutations that govern the Omicron S-mediated low fusogenicity and endosomal entry. We determined H655Y, and to lesser extent T547K, as the key mutation on the S protein of Omicron that critically governs its preferential endosomal entry route and impaired fusion activity, including for the recently emerged BA.4/5, BA.2.12.1 and BA.2.75 subvariants. Moreover, through molecular modelling, we provide evidence that these mutations likely stabilize the S trimer conformation by forming new molecular interactions. Together, these results provide insights for understanding the altered cellular tropism and pathogenicity for Omicron.

The most important finding of this study is that the altered entry route preference of Omicron is largely determined by the key H655Y mutations. It is well established that SARS-CoV-2 is capable of utilizing either endosomal entry mediated by Cat L/B or plasma membrane entry mediated by TMPRSS2 (Bestle *et al.*, 2020; Hoffmann *et al.*, 2020; Peacock *et al.*, 2021). However, SARS-CoV-2 entry in primary lung epithelial cells and lung-derived cell lines such as Calu-3 cell is largely TMPRSS2 dependent (Peacock *et al.*, 2021), likely occurring on the plasma membrane. Following its emergence, the Omicron variant BA.1 has been shown to have a distinct entry profile, utilizing predominantly the endosomal entry pathway (Pia and Rowland-Jones, 2022), exhibiting poor replication in lower airway derived primary cells and Calu-3 cells (Gupta, 2022), as well as displaying reduced disease severity (Halfmann *et al.*, 2022). We found that the H655Y mutation governs BA.1.1 entry through endosomes, as suggested by the significant increase in viral infectivity observed in HEK293T-ACE2 cells but substantial reduction in viral infectivity in Calu-3 cells, and by the increased sensitivity of H655Y bearing variants to E64d, yet with decreased sensitivity to Camostat, which are also supported by other recent studies (Hu et al., 2022; Yamamoto et al., 2022). However, the impact of H655Y on *in vivo* virus tropism and pathogenicity remains to be investigated. If the altered entry route preference introduced by the H655Y mutation is responsible for the enhanced nasopharynx tropism and reduced pathogenicity of the Omicron subvariants, any reversion of the H655Y mutation in future variants would be of great concern, as such a variant may exhibit enhanced pathogenicity. Careful monitoring this reversion mutation in the pandemic is warranted.

Membrane fusion is critical for entry of all enveloped viruses. We found that reversion mutations K547T and Y655H significantly promoted BA.1.1 S-mediated cell-cell fusion whereas forward mutations T547K and H655Y slightly impaired the D614G S-mediated cell-cell fusion, indicating that these two residues critically determine the low fusogenicity of BA.1.1. Quite unexpectedly, we found no evidence that T547K and H655Y affect S processing of BA.1.1 in cell lysates. Rather, reversion mutations K547T and Y655H strongly promote S1 shedding in the presence of sACE2, indicating that T547K and H655Y mutations, especially the latter, critically stabilize the BA.1.1 S conformation. This result is correlated with our structural modelling that T547K appears to stabilize the close conformation of S protein, which is also supported by a recent study reporting an extra hydrogen bond between the tyrosine residue at position 655 in S1 and the threonine residue at position 696 in S2 of BA.1 (Yamamoto *et al.*, 2022). Further structural analyses, including comparisons between K547T/Y655H reversion mutants and the parental BA.1.1 or other Omicron subvariants by cryogenic electron microscopy (cryo-EM) or crystallography, are needed to further elucidate the role of T547K and/or H655Y in Omicron subvariant S conformation.

It is important to note that H655Y mutation has been found to be associated with SARS-CoV-2 infection in index cats and minks (BraunID et al., 2021; Escalera *et al.*, 2022). In addition, the H655Y mutation appears to have arisen independently multiple times in human population, and is a lineage-defining mutation for the Gamma (P.1) SARS-CoV-2 variant in addition to the Omicron subvariants (Escalera *et al.*, 2022). Importantly, H655Y is present in all predominant Omicron sublineages, including BA.1.1, BA.1, BA.2, BA.2.12.1 and more recent BA.4/5 and BA.2.75, indicating that H655Y likely improves fitness and the ability to adapt to new hosts, including humans, cats, minks, and others. This is supported by a recent report demonstrating an enhancement of virus infectivity in mice for H655Y-containing viruses (Zhu et al., 2022).

Our findings in this work, along with other recent reports, together suggest that the occurrence of mutations at position 655 in S protein of current and future SARS-CoV-2 variants needs to be closely monitored. Additionally, *in vivo* examinations of the impact of the H655Y mutation on virus tropism and pathogenicity are critical and need to be investigated, as any reversion of the H655Y mutation could generate new concern for the course of the COVID-19 pandemic as novel Omicron subvariants continue to emerge.

## Acknowledgements

We thank the NIH AIDS Reagent Program and BEI Resources for supplying important reagents that made this work possible. S.-L.L. was supported by a fund provided by an anonymous private donor to OSU. J.P.E. was supported by Glenn Barber Fellowship from the Ohio State University College of Veterinary Medicine. K.X. was supported by the Ohio State University Comprehensive Cancer Center and a Path to K Grant through the Ohio State University Center for Clinical & Translational Science. The content is solely the responsibility of the authors and does not necessarily represent the official views of the university, or the Center for Clinical & Translational Science. P.-Y.S. was supported by NIH grants HHSN272201600013C and U01AI151801, and awards from the Sealy & Smith Foundation, the Kleberg Foundation, the John S. Dunn Foundation, the Amon G. Carter Foundation, the Summerfield Robert Foundation, and Edith and Robert Zinn.

## Author Contributions

S.-L.L. conceived and directed the project. P.Q. performed majority of the described experiments, and J.P.E. performed experiments related to infectious viruses. C.K. made the infectious Omicron BA.1 variants. C.Z. assisted in experiments. P.Q., J.P.E., and S.-L.L. wrote the paper. K.X. performed homology modeling. C.Z., Y.-M.Z., K.X., P.-Y.S., and X.X. provided insightful discussion and revision of the manuscript.

## Materials and Methods

### Cell Lines and Maintenance

HEK293T (ATCC CRL-11268, research resource identifier [RRID]: CVCL_1926), HEK293T-ACE2 (BEI NR-52511, RRID: CVCL_A7UK) and Vero E6 expressing high endogenous ACE2 (Vero-ACE2) (BEI NR-53726, RRID:CVCL_A7UJ) supplemented with 1% penicillin/streptomycin (MilliporeSigma, P4333) and 10% (v/v) fetal bovine serum (FBS) (Thermo Fisher Scientific, 26140-079). Calu-3, a gift of Estelle Cormet-Boyaka at The Ohio State University, were grown in Eagle’s Minimum Essential medium (EMEM) (ATCC, 30-2003), supplemented with 1% penicillin/streptomycin and 10% (vol/vol) FBS. HEK293T-ACE2 cells or Vero-ACE2 cells stably expressing TMPRSS2 were generated by transduction of pLX304 vectors expressing TMPRSS2 (a gift from Siyuan Ding at the Washington University in St. Louis), and selection by Blasticidine S hydrochloride (MilliporeSigma, 15205) (10 μg/ml for HEK293T-ACE2 cells and 7.5 μg/ml for Vero-ACE2 cells) for 7-14 days. All cell lines in this study were maintained at 37°C in the presence of 5% CO_2_.

### Plasmids and constructs

Constructs harboring different mutations were generated by long PCR mutagenesis based on pcDNA3.1-SARS-CoV-2-Flag-S-Flag-D614G, pcDNA3.1-SARS-CoV-2-Flag-S-Flag-BA.1.1, pcDNA3.1-SARS-CoV-2-Flag-S-Flag-BA.1, pcDNA3.1-SARS-CoV-2-Flag-S-Flag-BA.2, pcDNA3.1-SARS-CoV-2-Flag-S-Flag-BA.2.12.1, pcDNA3.1-SARS-CoV-2-Flag-S-Flag-BA.4/5 or pcDNA3.1-SARS-CoV-2-Flag-S-Flag-BA.2.75. HIV-1 NL4.3-inGluc was from Marc Johnson at the University of Missouri (Columbia, Missouri, USA).

### Generation of BA.1 FL-mNG and related mutant viruses

Infectious cDNA clone of USA-WA1/2020 and Omicron (BA.1) SARS-CoV-2 were generated using PCR-based mutagenesis as previously described (Pitts et al., 2022; Xie et al., 2020). To construct spike mutant K547T or Y655H mutant viruses, nucleotide substitutions were introduced through standard mutagenesis approach into a subclone pcc1-CoV-2-BA.1-F567 containing the spike gene of SARS-CoV-2 BA.1. The full-length infectious cDNA clone of SARS-CoV-2 was assembled by in vitro ligation of three contiguous cDNA fragments following the previously described protocol (Liu *et al.*, 2022b; Xie *et al.*, 2020). In vitro transcription was performed to synthesize full length viral genomic RNA. The RNA transcripts were electroporated in Vero E6 cells expressing TMPRSS2 (purchased from SEKISUI XenoTech, LLC) to recover the mutant viruses. Viruses were rescued post 2-4 days after electroporation and served as P0 stock. P0 stock was further passaged once on Vero E6 cells expressing TMPRSS2 to produce P1 stock. Spike gene was sequenced from all P1 stock viruses to validate the mutations. P1 stock was titrated by plaque assay on Vero E6 cells expressing TMPRSS2 and used for subsequent experiments. All virus preparation were carried out at biosafety level 3 (BSL-3) facility at the University of Texas Medical Branch at Galveston.

### Virus production and infection

HEK293T cells were transfected with 0.7 μg of different Spike plasmids along with 1.4 μg of HIV-1-NL4.3-inGluc at a 1:2 ratio using polyelthylenimine transfection (Cui et al.). Then, 48-72 hrs post-transfection, the culture supernatants were harvested. After removing the cell debris by spinning down at 3000 g for 10 minutes, the viruses were aliquoted and stored at −80°C. Pseudovirions were transduced into various cell lines; 6 hrs post-transduction, the media were changed. Gaussia luciferase activity was measured at 48-96 hrs after infection to determine the relative infectivity or entry efficiency of the indicated viruses.

For the inhibition assay using pseudotyped viruses, HEK293T-ACE2-TMPRSS2 cells were pretreated with indicated concentrations of EST/E-64D (E64d) (Sigma, 330005) or camostat mesylate (Camostat) (Sigma, SML0057) for 1 h, followed by transduction with pseudovirions of interest in the presence of the same concentrations of the drugs. After changing the media 6 hpi, the luciferase activity was measured at 48 and 72 hpi, respectively.

For the inhibition assay using infectious viruses, Vero-ACE2-TMPRSS2 cells were pretreated with indicated concentrations of EST/E-64D (E64d) (Sigma, 330005) or camostat mesylate (Camostat) (Sigma, SML0057) for 1 h, followed by infection with infectious viruses in the presence of the same concentrations of the inhibitors for 24 hrs. Subsequently, the cells were fixed with 3.7% formaldehyde for 1 h and analyzed by a Life Technologies Attune NxT flow cytometer.

### Flow cytometry

HEK293T cells used for the production of virions were washed with 1×PBS once, detached with 5 mM EDTA in PBS, washed with washing buffer (1×PBS containing 2% FBS) twice, following by fixing with 3.7% formaldehyde for 10 min. Then the cells were incubated with anti-SARS-CoV-2 Spike antibodies (T62) for 1 hr on ice. After three washes with cold washing buffer, the cells were incubated with FITC-conjugated anti-rabbit IgG antibodies for 1 hr. Subsequently, the cells were washed twice and analyzed by a Life Technologies Attune NxT flow cytometer.

### Syncytia formation assay

HEK293T-ACE2 cells were cotransfected with GFP constructs and parental D614G S, Omicron S, or their related mutants, followed by imaging the syncytia formation under a Leica DMi8 fluorescence microscope after 24 h of transfection. The cell-cell fusion efficiency was analyzed by measuring areas of the fused cells from three representative images using the “Leica Application Suit X” software. Scale bars represent 150 μm.

### Western blotting

Western blotting was conducted as previously described (Zeng *et al.*, 2020). Briefly, cells were collected and lysed in 200 ul of RIPA buffer (50 mM Tris (pH 7.5), 150 mM NaCl, 1 mM EDTA, Nonidet P-40, 0.1% SDS) in the presence of protease inhibitor cocktail (MilliporeSigma, P8340), followed by clarification at 13200 rpm for 10 minutes, and boiling for 10 minutes at 100°C with 1x SDS loading buffer. To determine the Spike content in virion particles, pseudovirus supernatant was collected, filtered and purified by ultracentrifugation through a 20% sucrose cushion. The purified virions were dissolved in 1x SDS loading buffer. Subsequently, the samples were separated by 10% SDS-PAGE gels, transferred to PVDF membranes and immunoblotted with anti-S1 (Sino Biological, 40150-T62), anti-S2 (Sino Biological, 40590-T62), anit-GAPDH (Santa Cruz, sc-47724), and anti-p24 (anti-p24 (NIH ARP-1513) antibodies, followed by immunoblotting with anti-mouse-IgG-Peroxidase (Sigma, A5278) or anti-rabbit-IgG-HRP (Sigma, A9169) antibodies.

### Soluble ACE2-induced S1 shedding assay

HEK293T cells were transfected with the indicated Omicron Spike constructs and were treated or untreated with soluble ACE2-Fc (sACE2-Fc) at 37°C for 4 hrs. Subsequently, the cell culture media and cells were collected, and anti-FLAG beads (Sigma, F2426) were used to precipitate S1 subunit in the media. Following immune-precipitation, the samples were separated on 10% SDS-PAGE gel, and probed with anti-S1 (Sino Biological, 40150-T62), anti-S2 (Sino Biological, 40590-T62) and anti-GAPDH (Santa Cruz, sc-47724). Anti-mouse-IgG-Peroxidase (Sigma, A5278) and anti-rabbit-IgG-HRP (Sigma, A9169) were used as secondary antibodies.

### Structural modeling and analysis

Homology modeling of Omicron spike protein was performed on SWISS-MODEL server with cryo-EM structure of SARS-CoV2 G614 strain spike (PDB 7KRR) as template. The resulting homo-trimer spike structure has one RBD in up conformation and the other two RBD in down conformation. Residue examination and renumbering were carried out manually with program Coot. Inter-protomer interaction analysis was performed with PDBePISA server. Molecular contacts of Omicron mutants were examined and illustrated with the programs PyMOL.

### Statistical analysis

All experiments were conducted in at least three independent replications except for those specifically indicated. Statistical significance analyses were performed in GraphPad Prism 9 (San Diego, CA) and are referenced in the figure legends and include one-way Analysis of Variance (ANOVA) with Bonferroni’s post-tests to compute statistical significance between multiple groups for multiple comparison (**Figs. 1b-g**, **Figs. 3b, 3c, 3f, 3g, 3i, 3j** and **3l**, **Figs. S3b-c**, **Figs. S4b-c**, and **Figs. S4e-f**), and an unpaired, two-tailed Student’s t test with Welch’s correction was used (**Figs. 4a**, **4b**, and **4d**).

