## Supplemental Figures 1-4 for "Determinants and Mechanisms of the Low Fusogenicity and Endosomal Entry of Omicron Subvariants"

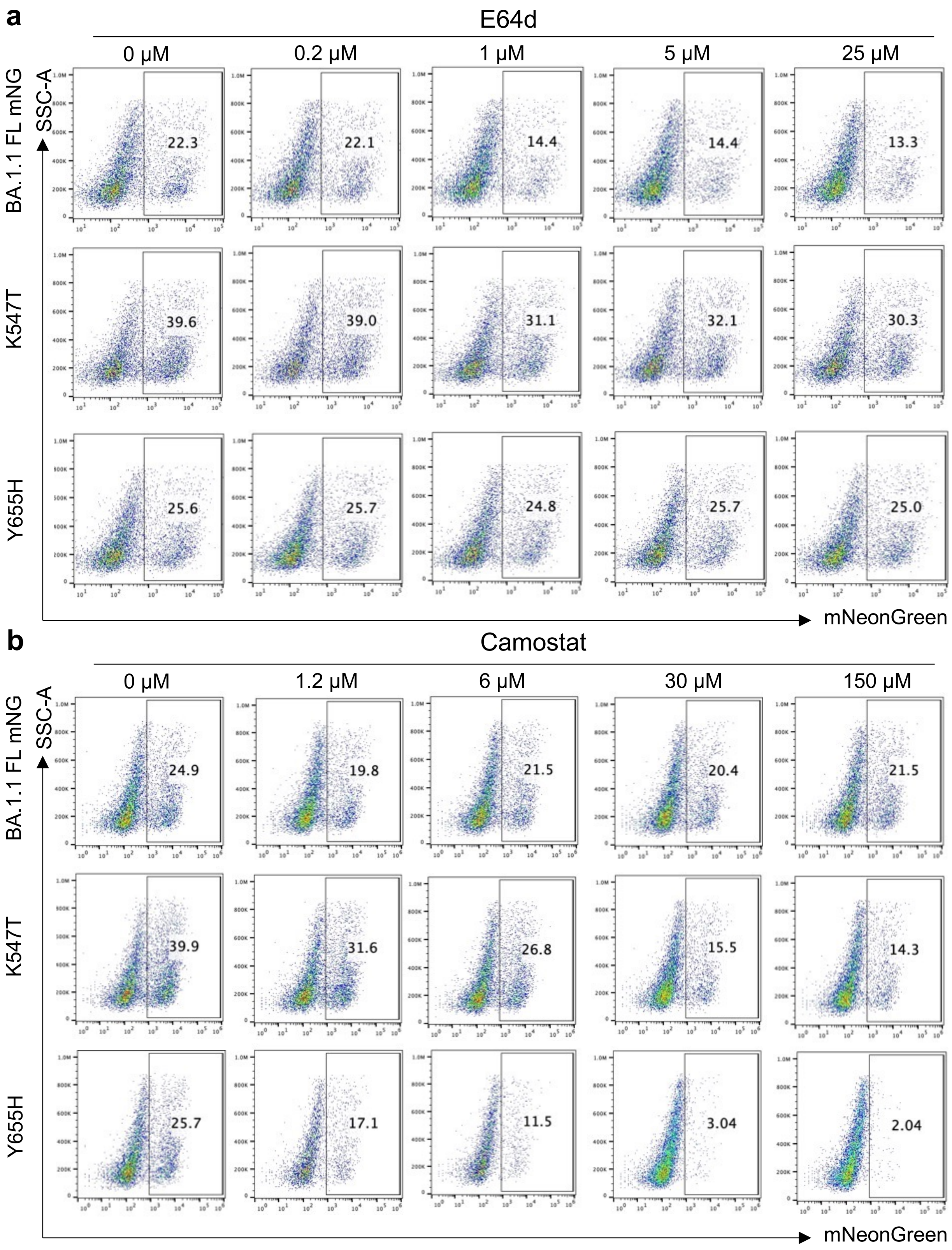


**Figure S1. H655Y dictates infectious BA.1.1 infection through the endosomal pathway**. Vero-ACE2-TMPRSS2 cells were pretreated with indicated concentrations of E64d (**a**) or Camostat (**b**), and then infected by the infectious SARS-CoV-2 viruses expressing mNeonGreen (Viana *et al.*). After fixing, the cells were analyzed by flow cytometry. Representative flow cytometry plots showing the percent of infection are presented.

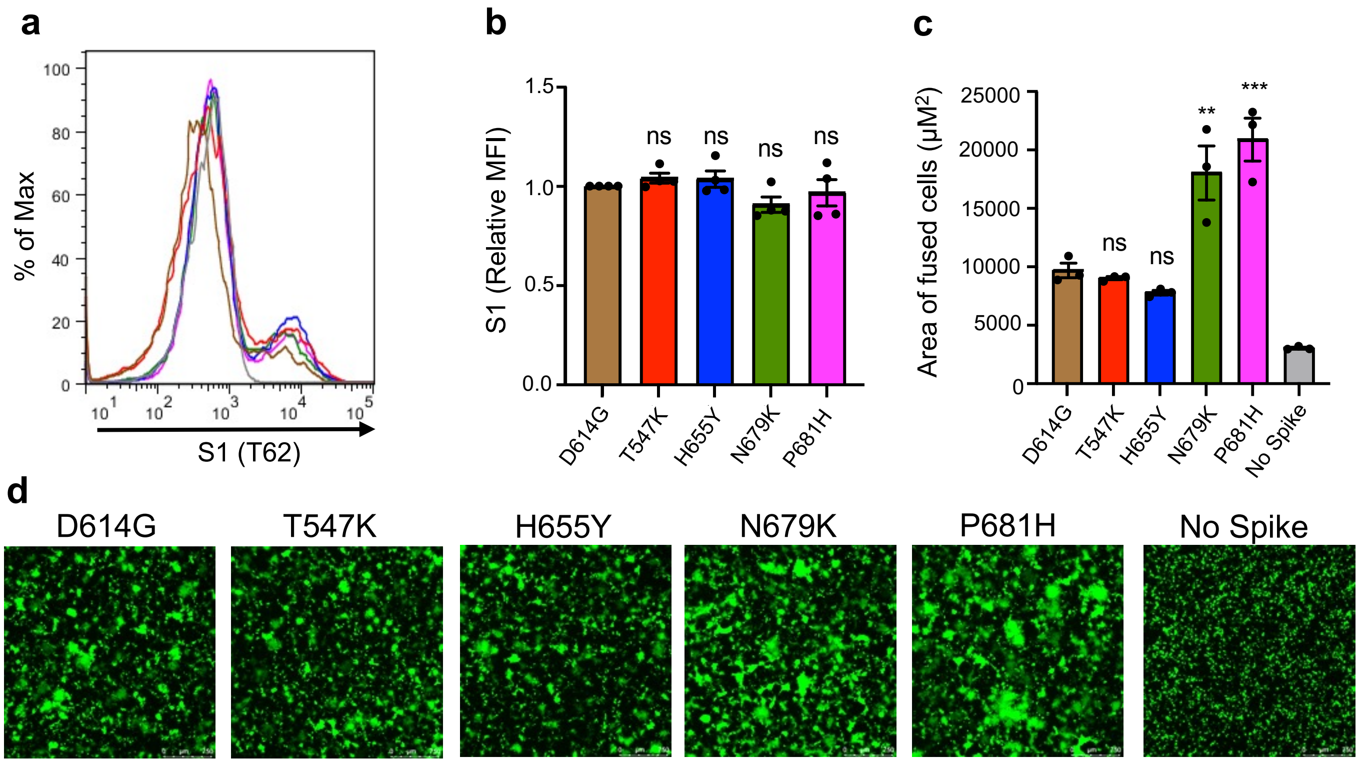


**Figure S2. H655Y mutation moderately impaired D614G S-mediated syncytia formation.** (**a-b**) Viral producer HEK293T cells were digested with EDTA/PBS and stained with anti-S1 (T62) antibody for flow cytometry. Representative flow cytometric analyses are shown in (**a**), and mean fluorescence intensity (MFI) is shown in (**b**), n=4. (**c-d**) HEK293T-ACE2 cells were transfected with the indicated S plasmids and a GFP expression plasmid, followed by fluorescence imaging and quantification of syncytia formation. Representative quantification (**c**) and images (**d**) are shown; n=3. Scale bars represent 150 μm. Error bars represent means +/- standard error. Significance was determined by one-way ANOVA with Bonferroni's multiple testing correction. ns indicates $p \geq 0.05$, $^{**}p < 0.01$, $^{***}p < 0.001$.

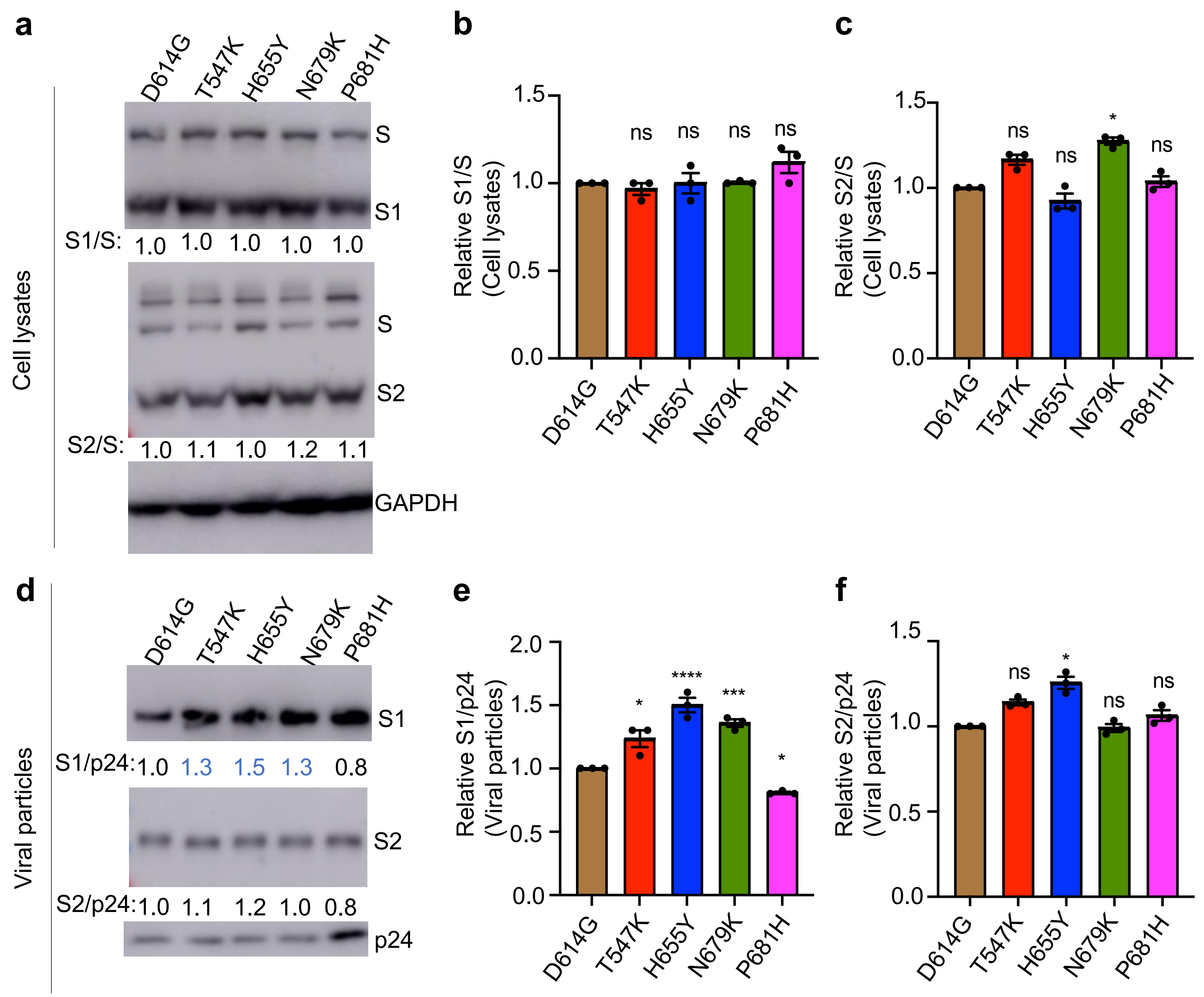


**Figure S3. Mutation H655Y contributes to more S incorporation into BA.1.1 virions.** Viral producer cells and supernatant were collected and blotted for S1, S2, GAPDH and p24 in cell lysates (**a**) and viral particles (**d**). The relative S1/S ratio in cell lysates (**b**), relative S2/S ratio in cell lysates (**c**), relative S1/p24 ratio in viral particles (**e**), and relative S2/p24 ratio in viral particles (**f**) are shown, n=3. Error bars are means +/- standard error. Significance was determined by one-way ANOVA with Bonferroni's multiple testing correction. P-values are represented as ns for $p \geq 0.05$, *$p < 0.05$, ***$p < 0.001$, ****$p<0.0001$.

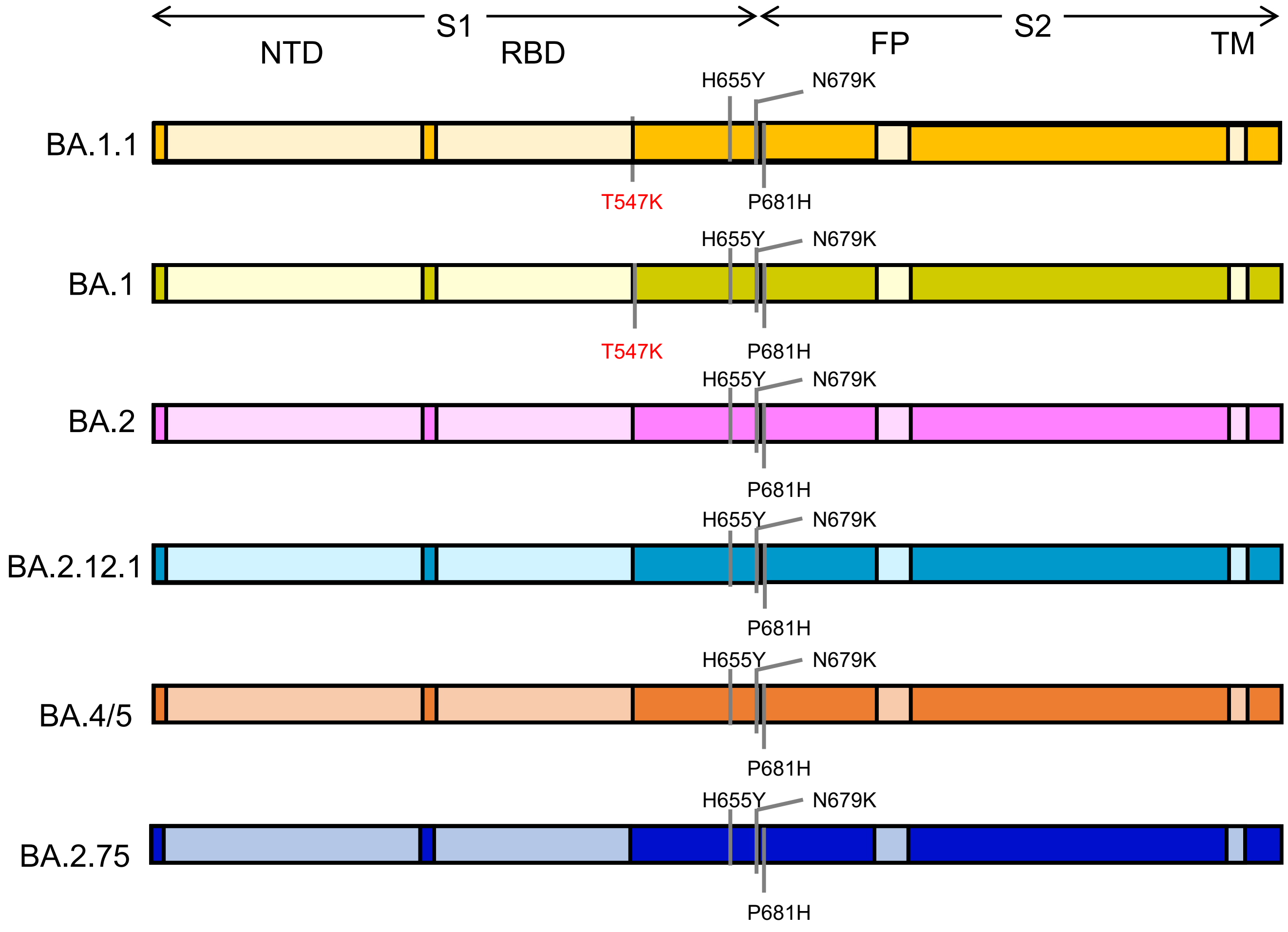


**Figure S4. Schematics representation of Omicron subvariants Spike glycoprotein**. Only the amino acid substitutions (relative to SARS-CoV-2 D614G S) at the C-terminus of S1, and/or near S1/S2 junction in BA.1.1, BA.1, BA.2, BA.2.12.1, BA.4/5 and BA.2.75 S are indicated. The T547K substitution, which is only included in BA.1.1 and BA.1 S, is highlighted in red.
